# An imaging framework for nuclei-based three-dimensional cell quantification in intact tissue using phase-contrast X-ray CT

**DOI:** 10.64898/2026.06.03.729871

**Authors:** T. Partridge, R. Ahmad, A. Astolfo, I. Buchanan, M. Endrizzi, M.A. Hawkins, A. Olivo, M. Esposito

## Abstract

**Objective:** Quantitative analysis of cellular morphology and spatial organisation within intact tissue remains challenging, particularly when three-dimensional information must be preserved. This study investigates whether laboratory propagation-based phase-contrast X-ray computed tomography (CT) can support nuclei-based quantitative analysis in unstained tissue while retaining the surrounding tissue architecture.

**Approach:** We propose a nuclei-based quantitative imaging workflow that combines laboratory propagation-based phase-contrast X-ray CT with volumetric segmentation. Intact unstained liver tissue was imaged at cellular resolution, then nuclei were segmented throughout the reconstructed volume and quantitative metrics describing the nuclear morphology and spatial organisation were extracted. The resulting measurements were evaluated in two non-overlapping volumes of interest and compared with histological reference data using thickness-matched virtual CT slices.

**Main results:** The proposed workflow enabled visualisation and segmentation of individual nuclei throughout intact unstained liver tissue. Comparable nuclear morphology and spatial organisation metrics were obtained from two non-overlapping volumes of interest, indicating stable segmentation performance within the analysed specimen. Comparison with H&E histology demonstrated broad agreement in nuclear morphology.

**Significance:** This work demonstrates the feasibility of nuclei-based quantitative analysis using laboratory phase-contrast X-ray CT in intact unstained tissue. The approach provides volumetric cellular information while preserving tissue architecture and may complement conventional histological assessment in applications requiring threedimensional analysis. These findings highlight the potential of laboratory phase-contrast CT for quantitative tissue characterisation and spatial cellular analysis.

## Introduction

Quantitative cellular analysis in intact three-dimensional tissues is increasingly important across biomedical research (Fei et al. 2022, Tian et al. 2021), where biological interpretation often depends not only on the number of cells present, but also on their morphology, spatial organisation, and relationship to the surrounding tissue architecture. Conventional approaches, however, remain largely restricted to two-dimensional techniques, including direct counting using Bürker chambers, indirect measurements such as total DNA quantification, and histological examination of thin tissue sections. As advances in biological and medical research drive increasing interest in three-dimensional cell cultures and complex tissue architectures, current methods are inadequate for capturing volumetric cellular organisation within extended specimens. Critically, no widely adopted standardised methodology currently exists for direct volumetric cell quantification in intact three-dimensional samples. The limitations of two-dimensional techniques are particularly evident in applications that require volumetric information or preservation of the specimen: these methods provide only local, non-volumetric measurements and are inherently destructive, preventing subsequent use of the sample (for example in tissue engineering workflows). Moreover, the preparation required for thin-section imaging can distort tissue morphology and complicate registration with prior imaging data (Savvidis et al. 2022). Non-destructive three-dimensional optical imaging approaches, such as two-photon (Xiao et al. 2023), three-photon microscopy (Yildirim et al. 2019) and micro-Optical Coherence Tomography (*µ*-OCT) (Yu et al. 2019), have demonstrated high-resolution volumetric imaging of tissue microstructure without the need for tissue clearing and provide valuable complementary tools for three dimensional tissue characterisation. However, these techniques remain subject to trade-offs in imaging depth, field of view and contrast in thick biological specimens. More generally, optical methods become increasingly constrained as sample thickness increases. Light scattering in thick biological specimens severely limits imaging depth, while tissue clearing procedures, although capable of improving optical penetration and enabling threedimensional visualisation (Tian et al. 2021, Richardson et al. 2021, Yamada et al. 2026), are time consuming (up to several weeks for a 1-cm^3^ volume), disruptive to tissue integrity and often incompatible with downstream analyses (Brenna et al. 2022). Alternative optical methods such as z stack scanning (Fei et al. 2022) can generate three-dimensional information but suffer from anisotropic resolution, with slice thicknesses on the order of tens of micrometres, limiting their applicability for accurate single cell quantification.

X-ray Computed Tomography (CT) offers a promising route to overcoming these limitations by enabling non-destructive three-dimensional imaging of thick specimens. Owing to the substantially higher penetration power of X-rays relative to visible light, imaging is not constrained by tissue scattering, eliminating the need for harsh optical clearing procedures. Although conventional micro-CT can, in principle, provide micron scale spatial resolution suitable for single cell detection (Clark & Badea 2021, Tamminen et al. 2020, Palmroth et al. 2020), its suitability for cell quantification is hindered by the inherently low soft tissue contrast resulting from the weak attenuation differences between cells and their surrounding matrix (Esposito et al. 2026), unless heavy metal staining is employed. Conversely, X-ray phase-contrast imaging is particularly well suited to resolving subtle variations in soft tissue composition, as it exploits changes in the real part (*δ*) of complex refractive index (*n* = 1 − *δ* + *iβ*), rather than relying solely on attenuation contrast governed by *β*, which can be several orders of magnitude lower than *δ* in the hard X-ray regime. Extensive demonstration of the potential of X-ray phase-contrast for cell-scale resolution in tissue sample has been demonstrated in the synchrotron environment (Eckermann et al. 2021, Bosch et al. 2022, John et al. 2026). Translating these capabilities to laboratory systems remains challenging, as current benchtop approaches seldom achieve reliable cellular resolution in unstained tissue (Esposito et al. 2026, Topperwien et al. 2018).

Here, we introduce an integrated quantitative imaging framework in which laboratory phase-contrast X-ray CT is combined with volumetric segmentation and histological comparison for nuclei-based analysis in intact unstained tissue. The approach enables individual nuclei to be visualised throughout the imaged tissue volumes and allows metrics describing nuclear morphology and spatial organisation to be extracted, while retaining surrounding tissue architecture. Importantly, we show that this workflow can be implemented using the simplest form of X-ray phase-contrast imaging, namely propagation-based imaging (see Methods). The combination of micron scale isotropic spatial resolution (≈1 *µ*m, for the microscope used in this study, as previously shown (Esposito 2025)) and the enhanced soft-tissue contrast arising from phase effects enables individual nuclei to be resolved in unstained tissue. Using ex vivo human liver tissue as a model system, we applied the workflow to two spatially distinct, non-overlapping volumes of interest from the same specimen and examined the consistency of the extracted measurements within the imaged tissue core. Comparison with H&E histology using thickness-matched virtual CT sections showed substantial overlap and broad comparability between the nuclear morphology distributions. This work show the feasibility of laboratory phase-contrast X-ray CT for nuclei-based quantitative analysis in three dimensions and support its potential use as a complementary approach for investigating cellular organisation within the context of volumetric tissue architecture.

## Methods

### Tissue specimens

Ex vivo human liver tissue was used in this study. Tissue was obtained from two resection specimens following diagnosis of liver metastasis secondary to colorectal cancer, under Royal Free London RFL Biobank Ethical Reference number NC2021.20 and National Research Ethics Service (NRES) Research Ethics Committee (REC) number 21/WA/0388. Specimens were fixed in 4% PFA (paraformaldehyde), dehydrated in ethanol through increasing concentrations (70–100%), and imaged in ethanol. Imaging in 100% ethanol was chosen, as previous work (Maughan Jones et al. 2026) showed that, for phase contrast X-ray imaging, increasing ethanol concentration leads to improved soft tissue contrast compared with lower ethanol concentrations and, more generally, compared with samples immersed in aqueous solutions.

### Sample preparation

A combination of planar and tomographic imaging was used. For planar imaging, specimens were cut into 250 *µ*m-thick slices (Leica VT1000S). For CT, a 1-mm-diameter biopsy punch was taken from a larger wet specimen at the fixation stage. Planar sections were mounted on thin Kapton foils, and the CT core was mounted in a 1-mm-diameter Kapton tube and sealed with wax. After imaging, the 1-mm-diameter tissue core was embedded in paraffin, sectioned in 5 *µ*m thick slices, and stained with haematoxylin and eosin (H&E).

### X-ray microscope

Imaging was performed using a free-space propagation-based phase-contrast X-ray imaging system. A rotating-anode X-ray source (Rigaku RA-Micro7 HF) with a copper anode was coupled to a doubly bent multilayer optic to select the Cu K*α* lines and focus the beam to a ∼350 *µ*m focal spot. An indirect-conversion detector was used, comprising a scintillator, objective, tube lens and a CMOS sensor (Photometrics Prime 95B). The effective pixel size was 1.1 *µ*m, corresponding to an optical magnification of 10*×*. The experimental set-up has been described previously (Esposito 2025), and a schematic is shown in Figure 2.

**Figure 1.**
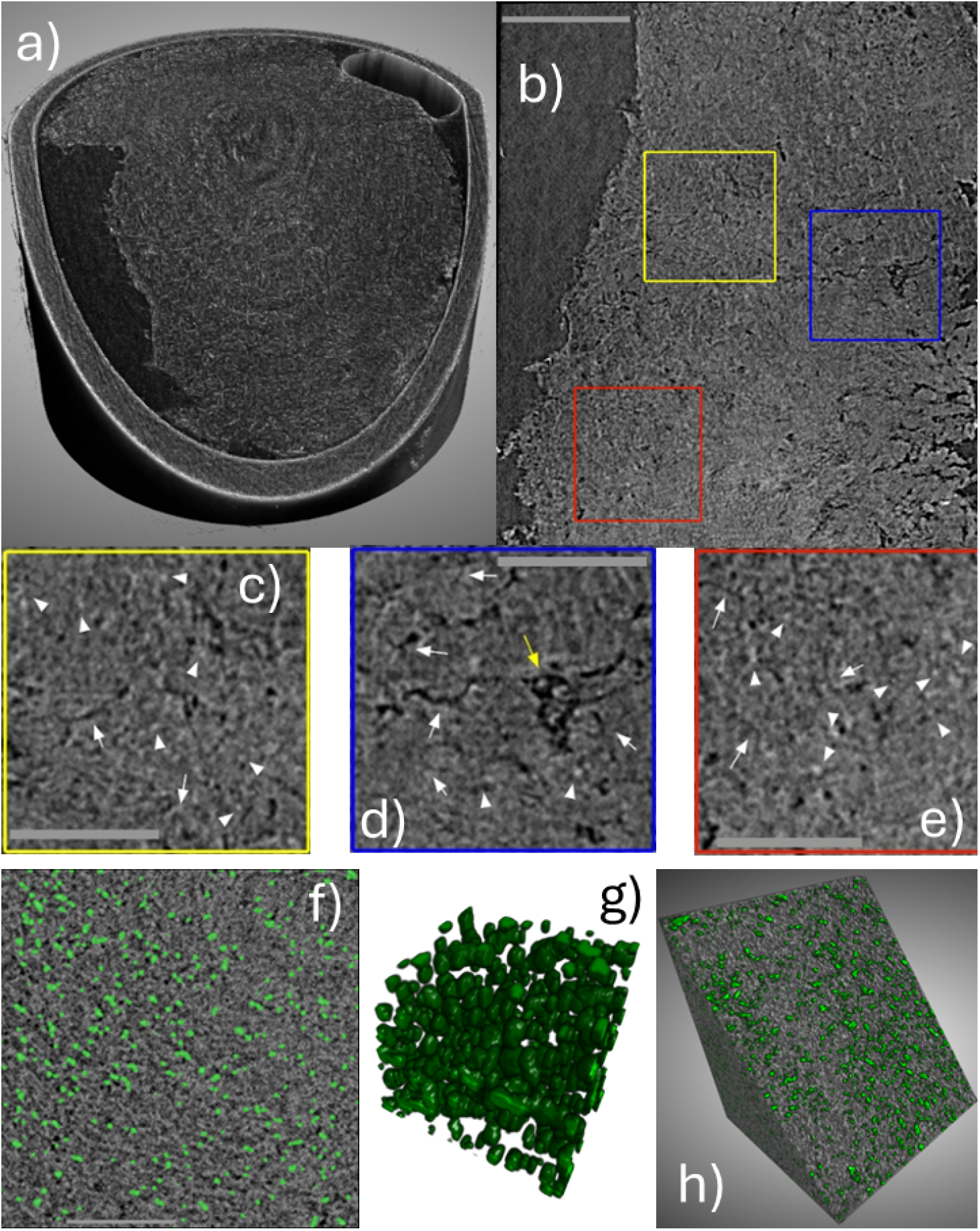
Imaging of liver tissue microstructure and nuclear segmentation. (a) Three-dimensional rendering of the liver tissue volume with a non-orthogonal cut-out plane showing tissue architecture, together with the Kapton tube walls, ethanol and an air bubble. (b) Sagittal slice illustrating the range of microstructural features visible in the phase-contrast CT data (scale bar, 200 *µ*m). Regions of interest are indicated by yellow, blue and red boxes in (b), with corresponding magnified views shown in (c), (d) and (e), respectively. Large vessels, small vessels and cell nuclei are indicated by yellow arrows, white arrows and white arrowheads, respectively (scale bars, 100 *µ*m). (f) Example CT slice region (330 *×* 330 *µ*m^2^) with segmented nuclei overlaid in green (scale bar, 100 *µ*m). (g) Three-dimensional visualisation of segmented nuclei. (h) Three-dimensional rendering of a volume of interest (330 *×* 330 *×* 330 *µ*m^3^) with an oblique cut-out plane (i.e. not orthogonal to the reference axes) and segmented nuclei overlaid in green. This volume, referred in the remainder of the manuscript as VOI 1, was chosen away from the container edges and from any reconstruction artefacts.

**Figure 2.**
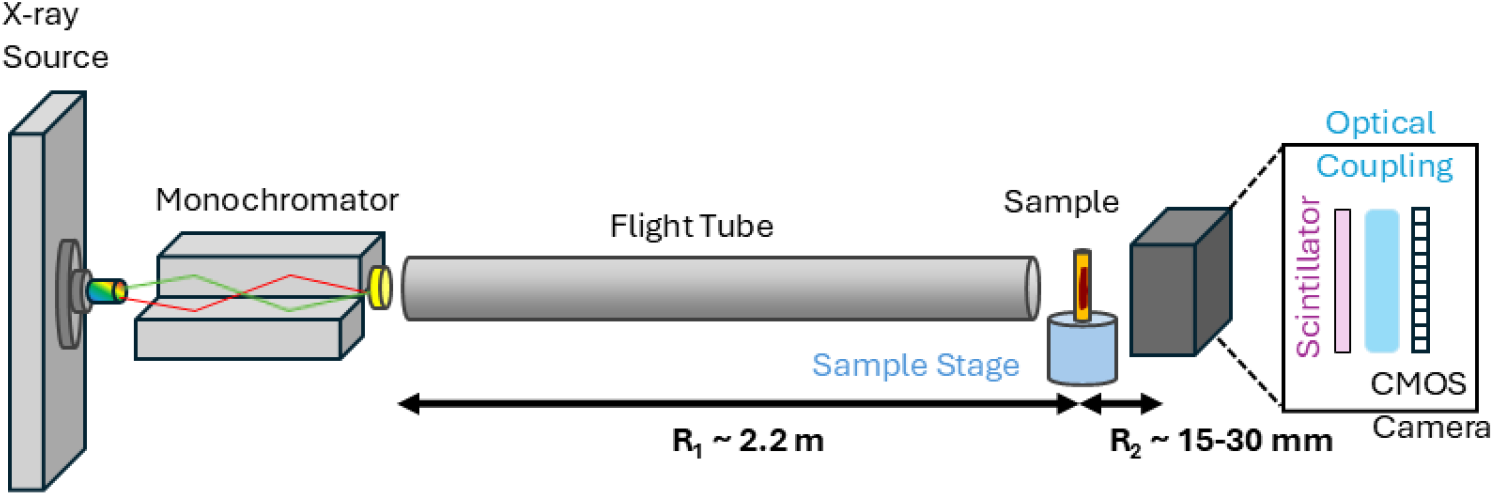
Experimental set-up of the laboratory propagation-based phase-contrast X-ray microscope. The sample to detector distance (*R*_2_) is adjustable, enabling control of the propagation distance.

Propagation-based phase contrast (Wilkins et al. 1996) relies on introducing a propagation distance between the sample and detector such that sample-induced phase shifts are converted into intensity variations at the detector plane. In this study, the sample was positioned approximately 2.2 m from the source and the sample-to-detector distance was set to 18 mm. The rationale for this propagation distance is discussed in the next section.

### Optimisation of the imaging set-up

To identify the optimal propagation distance balancing contrast, spatial resolution and noise performance, planar scans were acquired at four distances (15, 20, 25 and 30 mm). Planar measurements were acquired with a 10 s exposure time, with 50 frames collected and averaged. Representative liver tissue images obtained at 30 mm are shown in Figure 3a, revealing both macroscopic structures, such as vessels, and finer microscale features. A magnified view of the highlighted region of interest (ROI) further illustrates the visibility of high density components (e.g., nuclei) and lower density or hollow structures corresponding to capillaries or micro vessels embedded within a more uniform parenchymal background. Nuclei (or clusters of nuclei) were segmented based on their intensity, edges and textures. Example images at the four distances with nuclei segmentation overlaid are shown in Figure 4. Planar scans capture the full thickness of the sample (250 *µm* in this case). As the typical nuclear diameter is approximately 5 *µm*, signals from multiple nuclei distributed along the X-ray propagation direction can overlap in the projected image. Consequently, some segmented regions can include clusters of overlapping nuclei rather than individual nuclear structures. which can cause overlapping structures and apparent merging of nuclei. The planar image analysis was therefore restricted to evaluating nuclear or nuclear-cluster visibility as a function of propagation distance. Nuclear morphological measurements were not extracted from these data.

**Figure 3.**
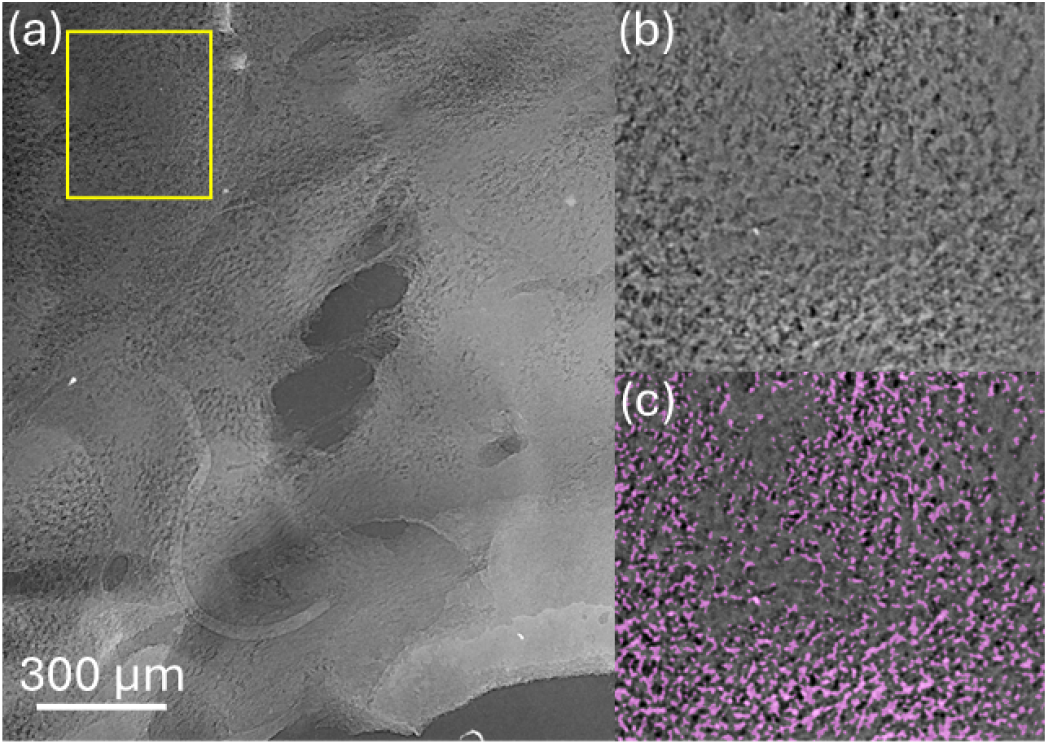
(a) Example planar scan of a liver specimen acquired at a 30 mm propagation distance; the yellow box indicates the region of interest (ROI). (b) Magnified view of the ROI. (c) Segmented nuclei overlaid on the magnified ROI. Please note that due to the sample thickness (250 *µm*) and to the 2D nature of the planar image, some regions can include clusters of overlapping nuclei rather than individual nuclei.

**Figure 4.**
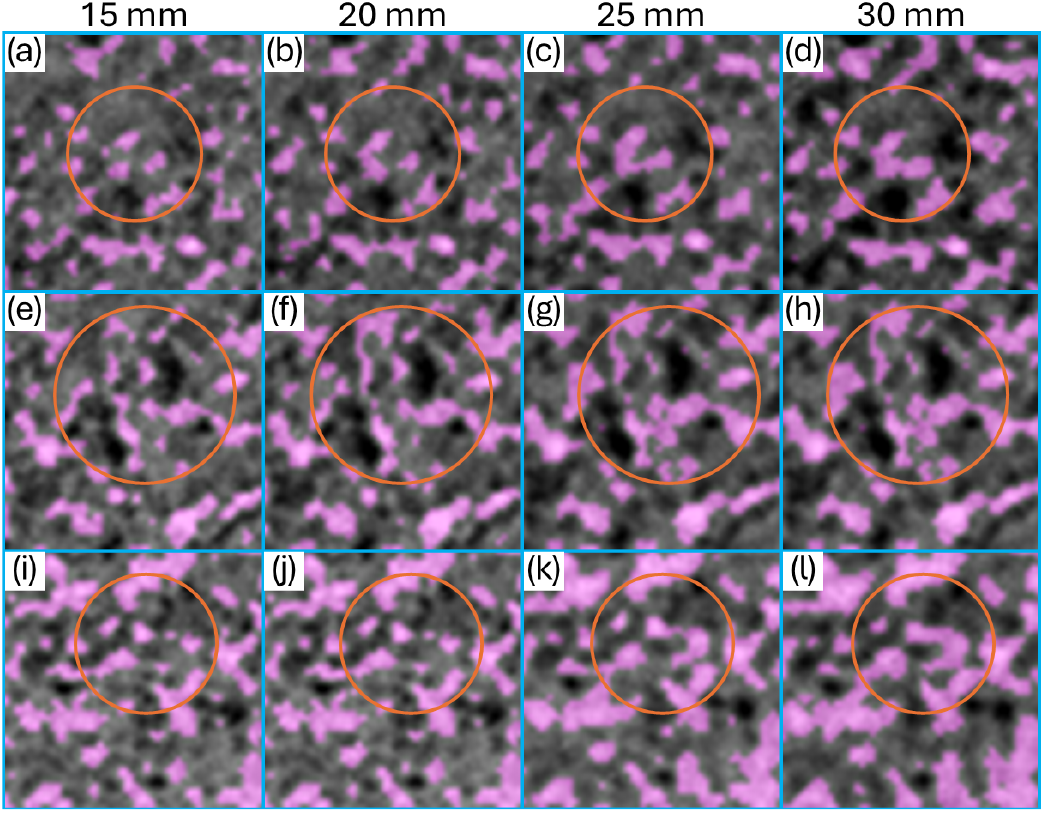
Segmentation results illustrate increasing amalgamation of nuclei with propagation distance as the pointspread function broadens. The three rows show different features with propagation distances of 15 mm for (a, e, i), 20 mm for (b, f, j), 25 mm for (c, g, k), and 30 mm for (d, h, l). Orange circles highlight regions where neighbouring nuclei begin to merge. All square boxes shown have a side dimension of 88 *µ*m.

Histograms of the selected nuclei and background are shown in Figure 5, illustrating clear separation of the nuclei against the remainder of the tissue. Additionally, standard image quality metrics, such as Signal to Noise Ratio (SNR) and Contrast to Noise Ratio (CNR) were calculated for nuclei relative to the surrounding tissue (background ROI, comprising pixels remaining after the exclusion of segmented nuclei and capillaries) to assess nuclei detectability as a function of the propagation distance. SNR and CNR are defined as follows:

**Figure 5.**
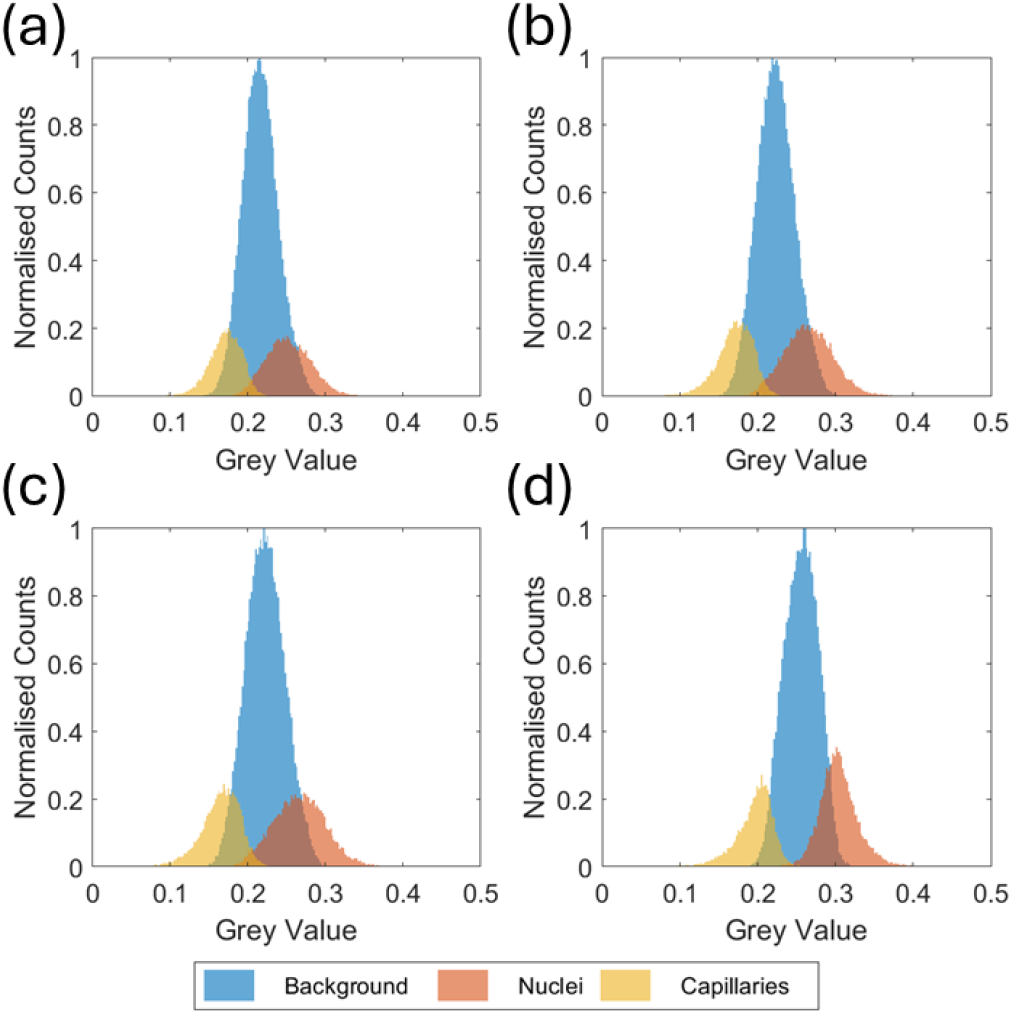
Histograms of segmented nuclei, capillaries and background (i.e., the remainder of the tissue), normalised to background counts for propagation distances of (a) 15 mm, (b) 20 mm, (c) 25 mm, and (d) 30 mm.

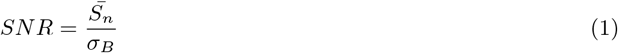

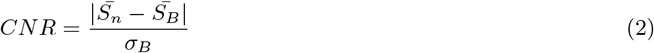

where 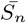 denotes the average signal measured for the segmented nuclei, and 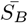 and *σ*_*B*_ represent the mean signal and standard deviation of the background (i.e. surrounding tissue excluding capillaries), respectively.

As reported in Table 1, both SNR and CNR reach their maximum values at a propagation distance of 30 mm. To further assess segmentation performance, nuclei were compared with their immediate surroundings by generating an external annulus via 2D Gaussian blurring of the segmented nuclei pixels. The width of the Gaussian blur was set to 3*σ* following a sensitivity analysis performed over the range 1.5*σ* to 5*σ*. The calculated CNR increased with increasing blur width up to approximately 2.5*σ*, beyond which it reached a plateau. This behaviour suggests that, for blur widths greater than 2.5*σ*, the annular region reliably samples non-nuclei pixels and yields stable background estimates. Consequently, a value of 3*σ* was selected for all subsequent analyses. Under this metric, CNR again peaked at 30 mm, whereas SNR was maximal at 15 mm (see Table 1).

**Table 1.**
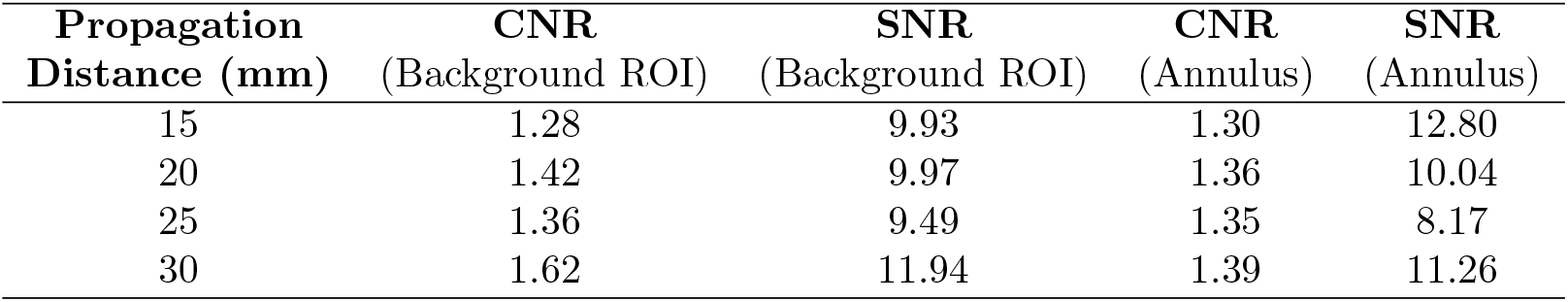
Contrast-to-noise ratio (CNR) and signal-to-noise ratio (SNR) for segmented nuclei, evaluated against the background ROI (i.e. comprising tissue pixels remaining after the exclusion of segmented nuclei and capillaries) and against an annulus representing the immediate surroundings of each nucleus.

| Propagation Distance (mm) | CNR (Background ROI) | SNR (Background ROI) | CNR (Annulus) | SNR (Annulus) |
| --- | --- | --- | --- | --- |
| 15 | 1.28 | 9.93 | 1.30 | 12.80 |
| 20 | 1.42 | 9.97 | 1.36 | 10.04 |
| 25 | 1.36 | 9.49 | 1.35 | 8.17 |
| 30 | 1.62 | 11.94 | 1.39 | 11.26 |

While CNR and SNR were used to assess nuclear contrast and detectability relative to the surrounding tissue, these metrics do not characterise spatial resolution. In propagation-based imaging, spatial resolution deteriorates with increasing propagation distance because of penumbral blurring. This dependence is well established theoretically (Gureyev et al. 2008) and has been characterised experimentally for the microscope used in this study in our previous work (Esposito 2025). In Figure 4, the effect of propagation-dependent blurring is apparent from the progressive merging of neighbouring nuclear signals with increasing propagation distance. In the corresponding segmentation overlays, nuclear regions that are separated at shorter distances become increasingly connected because blurring reduces the differences in intensity and texture at their boundaries, thereby decreasing segmentation reliability. Considering the SNR and CNR values reported in Table 1 alongside the qualitative evidence of blurring in Figure 4 and the previously characterised spatial resolution of the system, a propagation distance of 18 mm was selected as a pragmatic compromise between improved nuclear contrast and detectability and increased propagation-dependent blurring.

### Imaging and CT reconstruction

CT measurements were acquired with the same 10 s exposure time, collecting 2500 projections over 180^*◦*^ (angular step, 0.072^*◦*^). Each projection was acquired three times and averaged. The overall CT scan time was approximately 21 h. Sample drift and vibrations during acquisition were corrected by tracking a microsphere attached to the edge of the sample container and compensating for deviations from the ideal trajectories (Esposito 2025).

Phase retrieval using the Paganin method (Paganin et al. 2002) and tomographic reconstruction were performed in Python using TomoPy, with the filtered back projection (FBP) algorithm (Gürsoy et al. 2014).

### Segmentation workflow

Two-dimensional segmentation of planar X-ray and histology images was carried out in Ilastik (Berg et al. 2019) using intensity, edge and texture features. CT segmentation was performed in Avizo (Thermo Fisher Scientific) using texture classification based on first-order and histogram statistics as well as co-occurrence features with feature radius of 2 pixels and spherical texton of 3 pixels.

For the primary CT VOI (VOI 1), example nuclei and background tissue were manually seeded on the middle three slices, and the trained classifier was applied to the remainder of the sub-volume. Segmented nuclei were subsequently filtered by equivalent diameter to exclude objects outside the expected size range (2.4 *µ*m ≤ *D*_*eq*_ ≤ 11 *µ*m). Segmentation masks were exported and analysed in MATLAB using *bwconncomp* (3D connected components) and *regionprops* to extract nuclear metrics from the binary masks.

The secondary CT VOI (VOI 2) was seeded using the results from the first VOI (VOI 1), demonstrating the potential to reduce manual input. Avizo was selected for convenience in working with three-dimensional data; alternative trainable segmentation tools (e.g., Ilastik (Berg et al. 2019) or WEKA (Arganda-Carreras et al. 2017)) would be similarly applicable. Manual seeding was used in preference to deep-learning approaches (e.g., U-Net (Falk et al. 2019)) due to the limited size of the available training dataset.

Figure 6 summarises the segmentation workflow used in this study.

**Figure 6.**
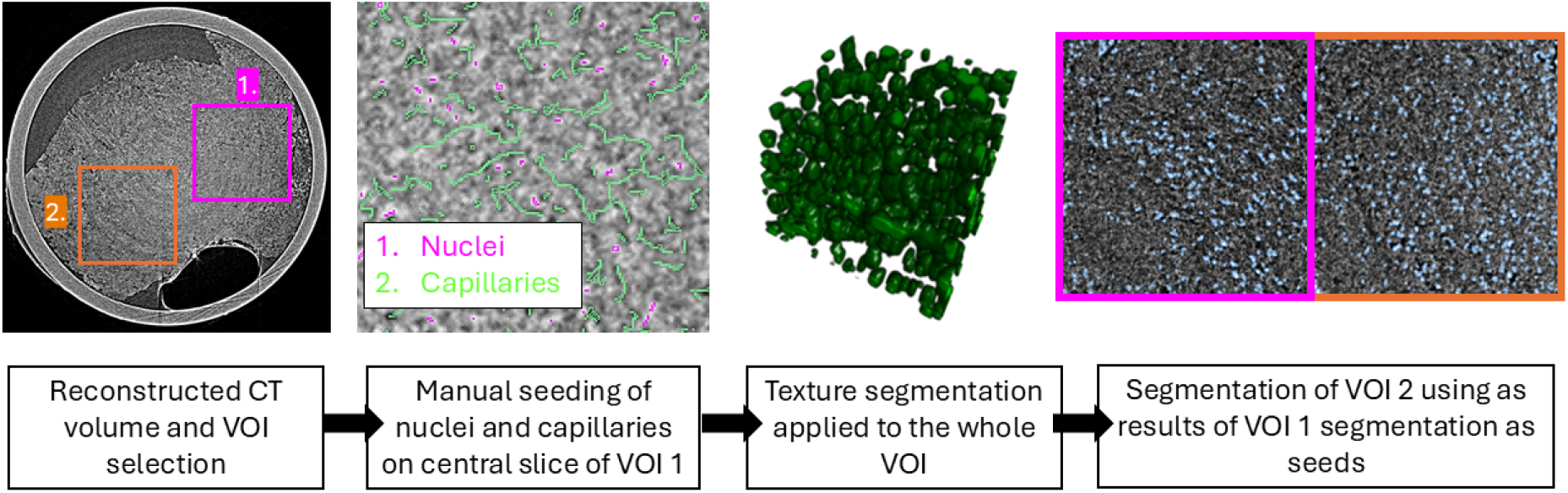
Segmentation workflow. A reconstructed phase-contrast tomogram is used to define volumes of interest (VOIs). Nuclei are segmented using Avizo texture classification after manual seeding on the middle three slices of VOI 1, producing 3D nuclear labels. The resulting classifier/labels are then used to initialise segmentation of VOI 2, illustrating the potential to reduce manual input when suitable training data are available.

### Quantitative analysis and histology validation

To assess the segmentation performance and consistency of measurements across regions, two non-overlapping volumes of interest (VOI 1 and VOI 2), each measuring 330 *µ*m *×* 330 *µ*m *×* 330 *µ*m, were selected at different heights within the reconstructed volume. Segmented nuclei were analysed to derive distributions of equivalent diameter, minor-to-major principal axes ratio, and three-dimensional nearest-neighbour distance. Mean values derived from these distributions were summarised as mean *±* standard deviation across all nuclei in each VOI.

Standard H&E histology was obtained to provide external validation of the segmentation and nuclei-based quantification workflow. An approximately matching region of the scanned liver sample, located away from the connective-tissue area, was selected for comparison. Exact spatial registration between CT and histology was not feasible. Therefore, representative regions with similar morphology were compared. To enable comparison with two-dimensional histology, oblique re-slices were extracted from each CT volume of interest. For consistency across modalities, each selected histology slice and CT re-slice was analysed over an identical in-plane field of view (330 *µ*m *×* 330 *µ*m). Nuclear metrics were computed from three slices (or CT re-slices) for each dataset and are presented on a slice resolved basis to visualise local sampling variability and slice-to-slice heterogeneity. Slice-based nuclear metrics included nuclear area, equivalent diameter, eccentricity and two-dimensional nearest neighbour distance. Using H&E histology as an independent reference, CT-derived metrics were interpreted in the context of the intrinsic slice-to-slice variability.

## Results

### Liver tissue imaging and nuclei-based cell quantification

A 1-mm diameter core, extracted from ex-vivo liver tissue, was imaged with the X-ray microscope described in the Methods section. As shown in Figure 1, the high-resolution phase-contrast CT data provide structural information across multiple spatial scales within the specimen. The three-dimensional rendering (Figure 1a) illustrates the overall volume and microstructural organisation. A sagittal slice (Figure 1b with magnified views in c-e) allows for the identification of different microstructural features, including vessels of varying size and cellular nuclei, which appear as high-density (bright) features. The clear delineation of cell nuclei in the threedimensional dataset provides sufficient contrast for robust segmentation using standard off-the-shelf tools. A description of the segmentation pipeline is provided in the Methods section. Figure 1f shows an exemplar CT slice in which segmented nuclei have been overlaid in green, while Figure 1g-h demonstrates the three-dimensional distribution of segmented nuclei within the tissue volume.

To assess segmentation performance and the consistency of the extracted measurements across spatially distinct regions, we selected two non-overlapping volumes of interest (VOI 1 and VOI 2), each measuring 330 *µ*m *×* 330 *µ*m *×* 330 *µ*m and positioned at different heights within the reconstructed volume. VOIs were not selected to represent different pathological regions or biological conditions. Rather, VOI 1 was used to establish the segmentation labels, which were subsequently applied to VOI 2 to examine whether the segmentation could be extended to a second region of the same tissue core with reduced additional manual input. Segmented nuclei were analysed as detailed in the Methods section, and the resulting distributions of equivalent diameter, minorto-major principal-axes ratio, and three-dimensional nearest-neighbour distance are shown in Figure 7. Mean values derived from these distributions are summarised in Table 2 (mean ± standard deviation (SD) across all nuclei in each VOI), demonstrating consistent measurements between VOIs across all parameters within the reported variability, compatible with sampling regions with similar tissue architecture.

**Table 2.** Quantitative 3D nuclear metrics for VOI 1 and VOI 2. The table reports the number of segmented nuclei and the mean *±* SD of equivalent diameter, principal axes ratio (minor/major; a proxy for nuclear sphericity), and 3D nearest-neighbour distance for all nuclei within each VOI.

| VOI | Number of Nuclei | Mean Equivalent Diameter ($\mu\text{m}$) | Principal axes ratio | 3D Nearest-Neighbour ($\mu\text{m}$) |
| --- | --- | --- | --- | --- |
| 1 | 19421 | $5.5 \pm 3.0$ | $0.43 \pm 0.11$ | $8.4 \pm 2.0$ |
| 2 | 20694 | $5.1 \pm 2.9$ | $0.42 \pm 0.11$ | $8.5 \pm 2.1$ |

**Figure 7.**
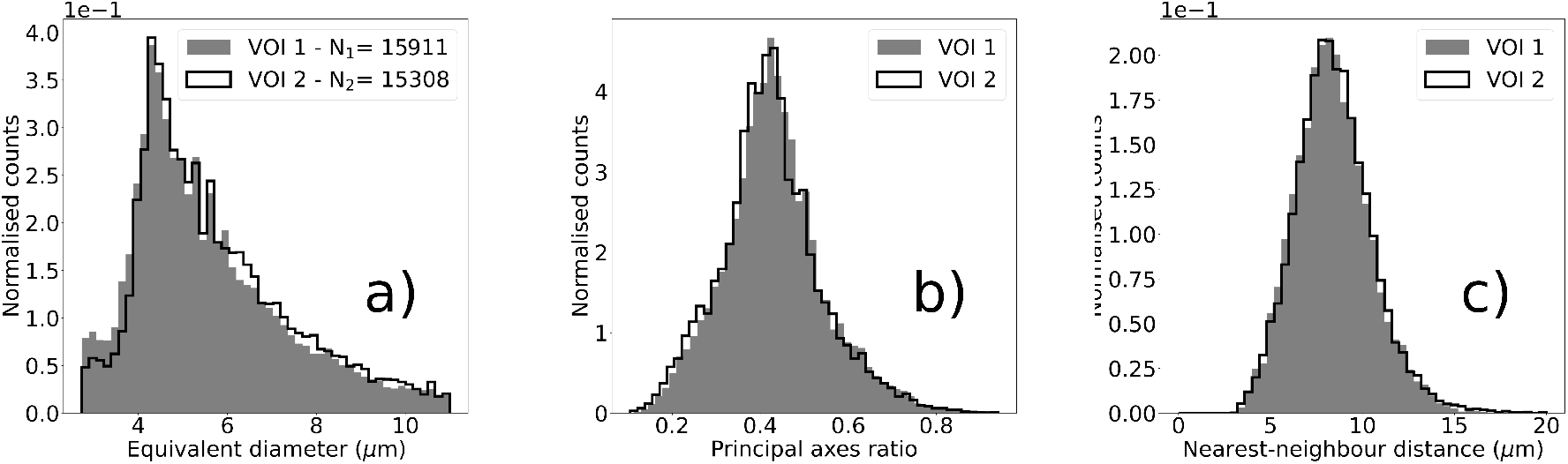
Distributions of 3D nuclear metrics across two volumes of interest. Histograms are normalised (probability density) and show the distributions of equivalent nuclear diameter (a), minor-to-major principal axis ratio (minor/major; sphericity proxy) (b), and three-dimensional nearest-neighbour distance (c) for nuclei segmented within CT VOI 1 and CT VOI 2. Summary statistics are reported in Table 1.

### Histological validation

Standard H&E histology was obtained to provide external validation of the segmentation and nuclei-based quantification pipeline. An example histological section from the scanned liver sample is shown in Figure 8a, displaying both hepatic and connective tissue characteristic of normal liver anatomy. A region approximately matching the one imaged with the laboratory system, and located away from the connective-tissue area, was selected for comparison (Figure 8b) and nuclei were segmented (Figure 8c). Exact spatial registration between CT and histology was not feasible because paraffin embedding and physical sectioning can introduce tissue deformation, shrinkage, changes in orientation and local tissue loss. Furthermore, the absence of fiducial markers or sufficiently distinctive internal landmarks prevented the precise position and orientation of each histological section from being identified within the CT volume. Representative regions with similar tissue morphology were therefore selected for comparison.

**Figure 8.**
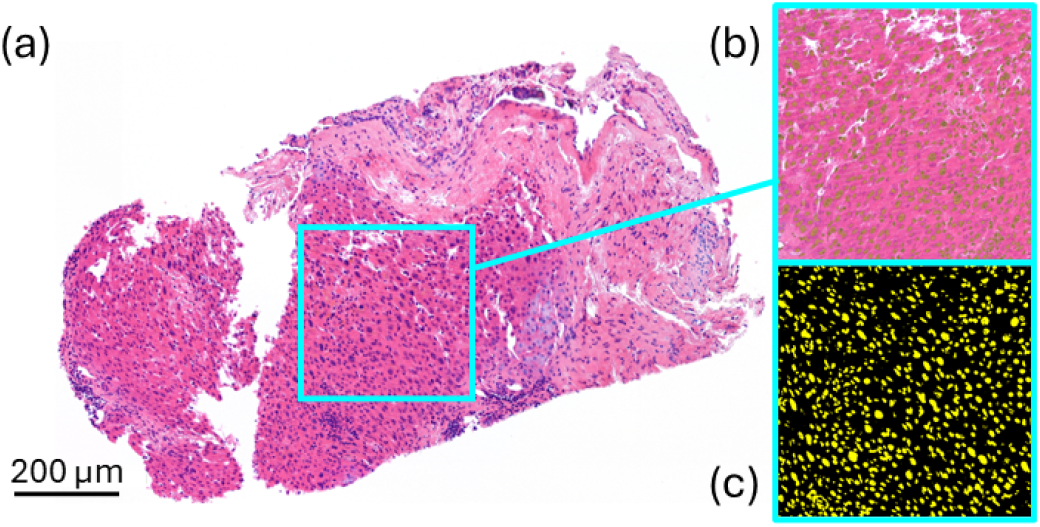
Histological reference for nuclear segmentation. (a) Representative H&E-stained histology section from the scanned liver sample. (b) Blue boxed region in (a) indicating the area selected for comparison, shown at higher magnification with the nuclear segmentation mask overlaid (green). (c) Nuclear segmentation mask for the same region shown separately.

To enable a direct comparison with the two-dimensional histology sections, virtual re-slices were generated from each CT VOI by extracting oblique arbitrary planes through the reconstructed volume. Subsequently, four consecutive slices were summed (binned), yielding an effective slice thickness of 4.4 *µm*. Re-slicing allowed visualising nuclei at locations within the volume that were independent of the detector orientation (rows and columns), while slice binning enabled the analysis of cellular morphology over a thickness comparable to that of the histological sections (≈5 *µm* thick). This approach facilitated a more direct comparison between CT-derived and histology-derived measurements by reducing the mismatch in section thickness between the two modalities.

Nucleus-level metrics are shown on a slice-resolved basis to capture local sampling variability within CT VOI 1, CT VOI 2 and H&E histology (Figure 9). For each dataset, metrics were computed from three slices (or CT re-slices), with each slice represented as a separate boxplot to visualise nucleus-level distributions and slice-to-slice heterogeneity for nuclear area, equivalent diameter, eccentricity and nearest-neighbour distance. Nearest-neighbour distances were computed within each two-dimensional slice (CT re-slice or histology section) using centroid-to-centroid distances. Dataset-level means and standard deviations, calculated across slices, are summarised in the top insets of each panel to provide an overview of slice-aggregated values and their dispersion. Using H&E histology as an independent reference, the CT-derived distributions showed substantial overlap and a broad comparability with histology-derived measurements. However, summary statistics for the two modalities do not appear to be quantitively consistent. The observed differences likely reflect a combination of spatial variability among sections and systematic modality-related effects, including section thickness, tissue preparation and segmentation.

**Figure 9.**
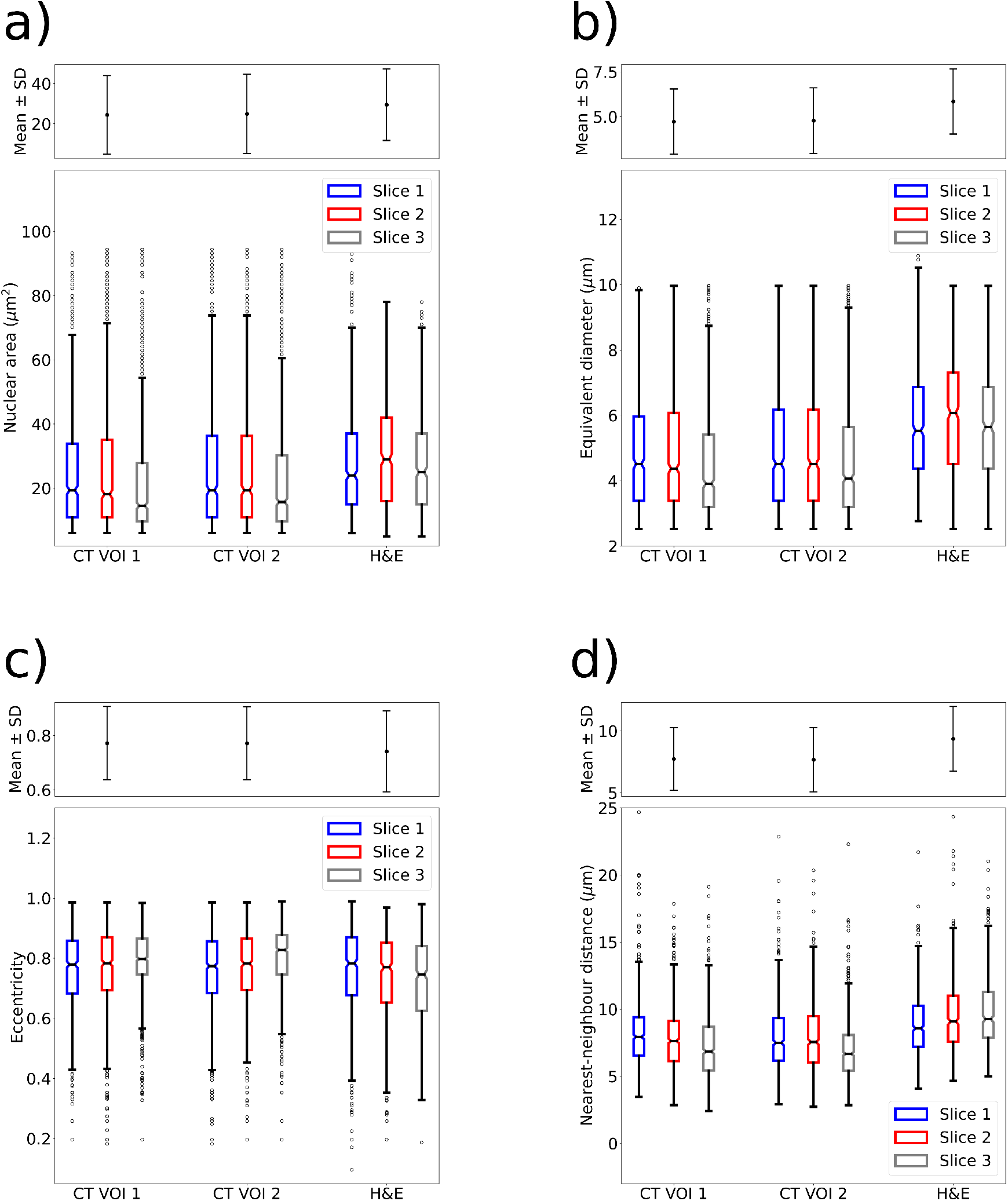
Nuclear metrics across CT and histology datasets. Nucleus-level distributions for individual slices (or re-slices) from CT VOI 1 (n=1569), CT VOI 2 (n=1518), and H&E histology (n=1671), shown as boxplots. Please note that each CT re-slice was obtained after binning four slices for a total thickness of 4.4 *µ*m. Here, *n* denotes the total number of segmented nuclei pooled across the three slices for each dataset. Distributions are shown for nuclear area (a), equivalent diameter (b), eccentricity (c) and nearest-neighbour distance (d). Nearest-neighbour distances were computed within each 2D slice using centroid-to-centroid distances. Boxplot elements: central mark indicates the median, and the bottom and top edges of the box indicate the 25^*th*^ and 75^*th*^ percentiles, whiskers extend to the most extreme data points, and outliers are shown individually; notches represent the 95% confidence interval of the median. Each boxplot corresponds to a single slice, explicitly visualising slice-to-slice heterogeneity within each dataset. Dataset-level means and standard deviations across slices are shown in the top insets in panels (a–d).

## Discussion

In this work, we demonstrate that combining a laboratory, propagation-based phase-contrast X-ray microscope with a robust segmentation workflow enables quantitative three-dimensional nuclear analysis in unstained, intact biological tissue, supporting nuclei-based estimation of cell density. In our approach, nuclear segmentation is used as a proxy for cell counting, consistent with common practice in optical microscopy where nuclear stains (e.g., DAPI) are routinely used and have been shown to provide an accurate estimate of cell density (McCaffrey et al. 1988). However, binucleation or multinucleation can limit the accuracy of nuclei-based cell quantification. This is particularly relevant for liver tissue, where hepatocytes can exhibit binucleation. Not all polyploid hepatocytes contain two distinct nuclei, and the proportion of truly binucleated cells is therefore expected to be lower than the proportion of polyploid hepatocytes (*<*25%)(Wilkinson & Duncan 2021). Where accurate cell counting is required, the impact of multinucleation can be addressed through appropriate correction strategies. Localised manual validation of true cell number in this dataset was not feasible because this would require delineation of individual cell membranes to determine whether one or more nuclei lie within each cellular boundary, whereas membranes were visible in neither the CT data nor standard histology. However, because polyploidy and associated multinucleation results in increased nuclear size (Carri`ere & Patterson 1962), future implementations of the counting workflow could incorporate nuclear area or three-dimensional volume, as shown in (Al-Khazraji et al. 2011), to correct nuclei-based estimates of cell number in case of multinucleation.

CT imaging was performed without slice sectioning, staining or optical clearing before image acquisition, while preserving the three-dimensional architecture of the extracted tissue core. The workflow therefore provides volumetric nuclear information that is complementary to conventional histology, potentially enabling CT data to guide the selection of regions and sectioning planes for subsequent histological analysis. This could support more targeted tissue sampling by identifying features or regions of interest within the three-dimensional volume before physical sectioning. Previous studies have shown that X-ray phase-contrast CT in the synchrotron environment, which typically involves higher radiation doses than those used in laboratory-based systems, does not introduce measurable structural or molecular alterations and is compatible with downstream histochemical, immunohistochemical, DNA, and RNA analysis (Li et al. 2023). This suggests that our pipeline has the potential to preserve compatibility with standard ancillary tests on the same specimen, allowing nuclei-based quantification to be paired with routine downstream analyses. We note, however, that the specific specimen used here exhibited drying artefacts, visible in histology, due to prolonged storage in 100% ethanol prior to embedding, rather than as a consequence of the X-ray imaging process itself. Future work will involve optimising the sample preparation method, particularly looking at imaging tissue in lower ethanol concentrations (70%), better suited for longer term storage, or directly as Formalin-Fixed Paraffin-Embedded (FFPE) which allows for long term storage and does not require any additional processing for validation histology to be completed. Additionally, the acquisition time in the present implementation remains relatively long and will be a focus of future optimisation. The flexibility of the imaging geometry provides scope to increase the detected photon flux, and thereby reduce exposure time, by shortening the source-to-sample distance (currently 2.2. m) and reducing the source size (e.g. by using different X-ray optics as shown in (Esposito et al. 2026)), while maintaining the spatial coherence required for phase-contrast imaging. Future work will evaluate these trade-offs to reduce scan duration while preserving sufficient spatial resolution and nuclear contrast for quantitative analysis.

The high-contrast and micron-scale spatial resolution of the imaging system allowed clear delineation of cell nuclei across large tissue volumes, enabling segmentation with standard, off-the-shelf tools. Importantly, analysis of two non-overlapping regions of interest revealed consistent metrics across all measured quantities, including equivalent diameter, nearest-neighbour distance, and sphericity. This demonstrates the stability of the pipeline across different spatial locations and microstructural contexts within the same sample.

Comparison with H&E histology provided an independent reference for assessing the CT-derived nuclear measurements. To reduce the systematic sampling mismatch between the two modalities, four consecutive CT slices were combined to produce virtual sections with an effective thickness of 4.4 *µm*, closely approximating the approximately 5 *µm* thickness of the physical histology sections. Following thickness matching, the CT-and histology-derived distributions showed substantial overlap and broad comparability. However, the H&E sections generally exhibited higher median nuclear areas and equivalent diameters than the CT re-slices, and the summary statistics were not quantitatively consistent between the two modalities. Although a large heterogeneity was observed for both modalities, differences in nuclear morphology may also reflect systematic modality-related effects, including the residual difference in section thickness, physical versus virtual section formation, tissue preparation and segmentation. Accordingly, this comparison supports the feasibility of extracting nuclear morphological information from laboratory phase-contrast CT, while highlighting the need for further validation, including direct spatial registration between CT and histology to enable more accurate cross-modality comparisons. The CT-derived measurements should instead be interpreted as complementary to histological measurements, with the additional advantage of enabling nuclear morphology and spatial organisation to be examined throughout a three-dimensional tissue volume.

There is growing interest in applying advanced segmentation and machine-learning approaches to enhance three-dimensional analysis and to support or replace existing 2D and manual cell-counting methods (Caicedo et al. 2017, Kataras et al. 2023). Generalisable segmentation frameworks, such as U-Net (Falk et al. 2019), offer particular promise for automated nuclei identification across diverse imaging conditions. In our framework, the first VOI (VOI 1) required manual seeding; however, the second VOI (VOI 2) was segmented entirely using the results generated from the first, without additional manual input. This demonstrates both the consistency of the segmentation procedure and the feasibility of developing a fully automated pipeline, such as a U-Net-based approach, given a broader dataset spanning multiple tissue types.

## Conclusion

This proof-of-principle study demonstrates the feasibility of using laboratory propagation-based phase-contrast X-ray CT to visualise and quantify individual nuclei in three dimensions within an unstained tissue core. The proposed workflow enables nuclear morphology and spatial organisation to be assessed throughout the imaged volume while preserving the surrounding tissue architecture. Laboratory phase-contrast CT may therefore provide a complementary approach to conventional histology by adding volumetric information on nuclear organisation. Future work will focus on validating the approach across additional specimens and tissue types, establishing direct CT-to-histology registration, optimising imaging geometry and sample preparation to reduce acquisition time, and developing more generalisable segmentation strategies, including deep-learning-based methods.

## Acknowledgments

This work was supported by the Medical Research Council (Grant No. MR/Y008448/1). Additional support was received from the EPSRC (Grants No. EP/T005408/1, No. EP/P023231/1, and No. EP/M028100/1). M. Esposito was supported by the NIHR Healthcare Engineering and Imaging Early Career Researcher Scheme. A.O. was supported by the Royal Academy of Engineering under the “Chairs in Emerging Technologies” scheme (CiET1819/2/78). RA and MAH were supported by the National Institute for Health and Care Research University College London Hospitals Biomedical Research Centre. Access to the Avizo software was provided by the National Research Facility for Lab X-ray CT (NXCT) through EPSRC grants EP/T02593X/1 and EP/V035932/1. We thank Dr J. C. Hutchinson (Great Ormond Street Hospital) for insightful interpretation of histological images.

## Notes

### Competing Interest Statement

The authors have declared no competing interest.

### Summary of Updates

Figure 9 updated to include slice binning

